# Surface microbiota of Mediterranean loggerhead sea turtles unravelled by 16S and 18S amplicon sequencing

**DOI:** 10.1101/2022.04.01.486708

**Authors:** Lucija Kanjer, Klara Filek, Maja Mucko, Roksana Majewska, Romana Gračan, Adriana Trotta, Aliki Panagopoulou, Marialaura Corrente, Antonio Di Bello, Sunčica Bosak

**Author notes:** **Correspondence:** Sunčica Bosak.

## Abstract

Loggerhead sea turtle is considered a keystone species with major ecological role in Mediterranean marine environment. As with other wild reptiles, their outer microbiome is rarely studied. Although there are several studies on sea turtle’s macro-epibionts and endo-microbiota, there has been little research on epibiotic microbiota associated with sea turtles’ skin and carapace. Therefore, we provide the first identification of combined epibiotic eukaryotic and prokaryotic microbiota on Mediterranean loggerhead sea turtles. In this study, we sampled skin and carapace of 26 loggerheads from the Mediterranean Sea during 2018 and 2019. To investigate the overall microbial diversity and composition, amplicon sequencing of 16S and 18S rRNA genes was performed. We found that the Mediterranean loggerhead sea turtle epibiotic microbiota is a reservoir of a vast variety of microbial species. Microbial communities between samples mostly varied by different locations and seas, while within prokaryotic communities’ significant difference was observed between sampled body sites (carapace vs. skin). In terms of relative abundance, Proteobacteria and Bacteroidota were the most represented phyla within prokaryotes, while Alveolata and Stramenopiles thrived among eukaryotes. This study, besides providing a first survey of microbial eukaryotes on loggerheads via metabarcoding, identifies fine differences within both prokaryotic and eukaryotic microbial communities that seem to reflect the host anatomy and habitat. Multi-domain epi-microbiome surveys provide additional layers of information that are complementary with previous morphological studies and enable better understanding of the biology and ecology of these vulnerable marine reptiles.

## 1 Introduction

Microbial communities associated with the external surfaces of animals represent an important part of the animal microbiome. The animal integument is a physical barrier between the internal environment that it protects, and the external environment that it interacts with. In vertebrates, one of the most extensively studied epimicrobiomes is the human skin (Turnbaugh et al., 2007; Byrd et al., 2018). The epidermal microbes enhance the skin barrier performance by modulating innate immunity and developing adaptive immunity (Sanford and Gallo, 2013), therefore helping to battle skin pathogens (Belkaid and Segre, 2014; Belkaid and Tamoutounour, 2016). A shift in the host’s health can alter the composition and functions of the skin microbiota that can lead to various diseases (Sanford and Gallo, 2013). Similarly, the native microbial communities of the epidermis can be affected by sub-optimal environmental conditions, negatively influencing their protective properties (Scharschmidt and Fischbach, 2013; Byrd et al., 2018).

The skin and other external body surfaces (e.g. horns, carapaces, hair and other keratinous hard tissues) of vertebrates differ between taxonomic groups and provide fairly diversified habitats for various animal-associated microbes (Ross et al., 2019). Since both intrinsic (e.g., species, sex, age) and extrinsic (e.g., geographic location, biotic and abiotic environmental conditions, captivity affecting the natural behaviour and diet) factors shape the community composition of the epimicrobiome, differences between even closely related host species or individuals are to be expected (Ross et al., 2019; Woodhams et al., 2020). Nevertheless, a certain degree of phylosymbiosis, a co-evolving microbiota harbouring the phylogenetic signal of their hosts (Brooks et al., 2016), is also observed (Ross et al., 2018). Apart from humans, much of the epimicrobiome research has focused on captive animals and pets as well as amphibians whose skin is more permeable and thus more susceptible to pollution and novel pathogens, potentially threatening the survival of entire populations and species (Ross et al., 2019). However, very little is known about the epimicrobiomes of reptiles, especially turtles, including terrestrial, freshwater, and marine species. Recently, there have been studies on external microbial communities of freshwater turtles (*Trachemys scripta, Pseudemys concinna* and *Emydura macquarii krefftii*) that identified major microbial components of eukaryotic and prokaryotic surface communities and showed that turtles’ microbiotas differ between body parts and between animals and their environment (McKnight et al., 2020; Parks et al., 2020). New knowledge about the functional and phylogenetic composition of epimicrobiomes of different species improves our understanding of the relationships between the host, its microbial flora, and the environment. Such advances in knowledge may contribute to a more efficient conservation of endangered and threatened macroorganisms. Skin microbiome research has a lot of potential in conservation biology of marine animals because of its accessibility and non-invasive sampling procedures. The potential of microbiome as bioindicator of ecosystem’s health has been recognized and effort is being put into the standardization of the methodology – e.g., from sampling to correct index calculations (Lau et al., 2015; Aylagas et al., 2017; Keeley et al., 2018; Cordier et al., 2019). It is possible that surface-associated microbiomes exhibit a stronger link with variations in the environment, while the internal microbial communities are more affected by the host’s intrinsic factors (Woodhams et al., 2020).

Although loggerheads are the most abundant sea turtle species in the Mediterranean Sea, they are threatened by coastal development, fishing bycatch, tourism, pollution and climate change (Casale et al., 2018). Skin and carapace of loggerheads provide habitats for a surprising variety of unique and taxonomically diverse macro-epibionts, including barnacles, amphipods and red algae (Hollenberg, 1971; Broderick et al., 2002; Frick and Pfaller, 2013). Some of these organisms require the sea turtle substratum to attach and thrive, and thus their survival is inextricably linked to the wellbeing and fitness of their hosts. The existing body of literature on loggerhead and other sea turtle microbiomes includes mainly studies investigating the internal microbiota, such as those living in the gut, cloaca, faeces, and oral cavities (Abdelrhman et al., 2016; Arizza et al., 2019; Biagi et al., 2019; Scheelings et al., 2020b, 2020a; Filek et al., 2021). The epimicrobiomes of sea turtles, in turn, have received far less attention. Recent years brought increased interest in micro-eukaryotic surface assemblages of sea turtles largely due to a series of projects exploring the diversity of sea turtle-associated diatoms (Majewska et al., 2015; Robinson et al., 2016; Rivera et al., 2018; Azari et al., 2020; Kanjer et al., 2020; Van de Vijver et al., 2020). Those studies identified a group of the diatom core taxa typical of sea turtles but also showed some biogeographic differences between diatom epizoic assemblages (Van de Vijver et al., 2020). Besides inventorial and ecological interest in diversity of epi-microbiome, there is a possible benefit for sea turtles’ health that could arise from these kinds of studies. For example, *Fusarium* spp. fungal infection of loggerhead eggs is considered a global threat (Bailey et al., 2018) and its detection on carapace and skin could be beneficial (Cafarchia et al., 2020). Further, evidence of antibiotic resistant bacteria found on sea turtles highlight the direct effect of antibiotic pollution in the seas (Pace et al., 2019; Alduina et al., 2020; Trotta et al., 2021). However, reports on prokaryotic and micro-eukaryotic non-diatom communities associated with the skin and carapace of sea turtles are extremely scarce and include a recent study by Blasi et al. (2022) that reported the composition of bacterial community based on 16S profiling from the carapaces of three juvenile loggerheads from the Tyrrhenian Sea.

In the present study we decided to further expand the perspective and study the whole microbial community found on external surfaces of loggerhead sea turtles from the Mediterranean Sea. According to the authors’ knowledge, this is the first report of the micro-eukaryotic diversity alongside the prokaryotic diversity using the 18S and 16S amplicon sequencing approach, respectively. Furthermore, we provide the first description of epi-microbial communities associated with the skin samples rather than just the loggerhead’s carapace. Detailed observation and statistical analyses of microbial assemblages’ taxonomic composition were addressed in accordance to our large and diverse loggerhead dataset. The role of many factors that could influence the microbial communities is considered and additional approaches in studies of this type are discussed.

## 2 Materials and methods

### 2.1 Sampling

Twenty-six loggerhead sea turtles were sampled from four different Mediterranean areas: Adriatic (n=14), Ionian (n=9), Tyrrhenian (n=1) and Aegean Sea (n=2; Figure 1) following recommendations from Pinou et al. (2019). Two separate samples were collected from each turtle, one from the carapace and one from the skin. Biofilm scrapings were taken using clean toothbrushes and/or a sterilised scalpel and were resuspended in 96% ethanol in sterile 50 ml conical tubes immediately after collection. Carapace samples were collected randomly from an entire carapace, whereas skin samples were taken from the animal head, neck, and flippers. All samples were stored at -20 °C until further processing. One turtle (ID010) was sampled twice: immediately upon arrival to the rescue centre and after approx. one year in rehabilitation. In total, 54 samples were collected from August 2018 until November 2019 (Table 1). Due to the heterogeneity of sampled turtles, we differentiate a turtle’s origin localities from “sampling locality” (Table 1). Origin locality is the location where a turtle was found in the sea or on the beach and the sampling locality refers to the place where samples were obtained. Origin locality and sampling locality is identical for the turtles sampled where they were found but differs for the turtles that were being brought to rehabilitation centres. The turtles were sampled in three rescue centres (Marine Turtle Rescue Centre Aquarium Pula and Blue World Institute Lošinj in Croatia, and The Archelon Sea Turtle Protection Society in Greece) and one veterinary clinic (The Sea Turtle Clinic, STC, Department of Veterinary Medicine, University of Bari “Aldo Moro” in Bari, Italy). The sea turtle status was designated as “wild” if the turtle was sampled immediately after capturing without being immersed into the rehabilitation pool, and “admitted” if the animal was admitted to a rehabilitation centre and was immersed in the rehabilitation pool prior to sampling. Time between the turtle admission and the sampling of its biofilm spans between one and ten days (except for ID010).

**Figure 1.**
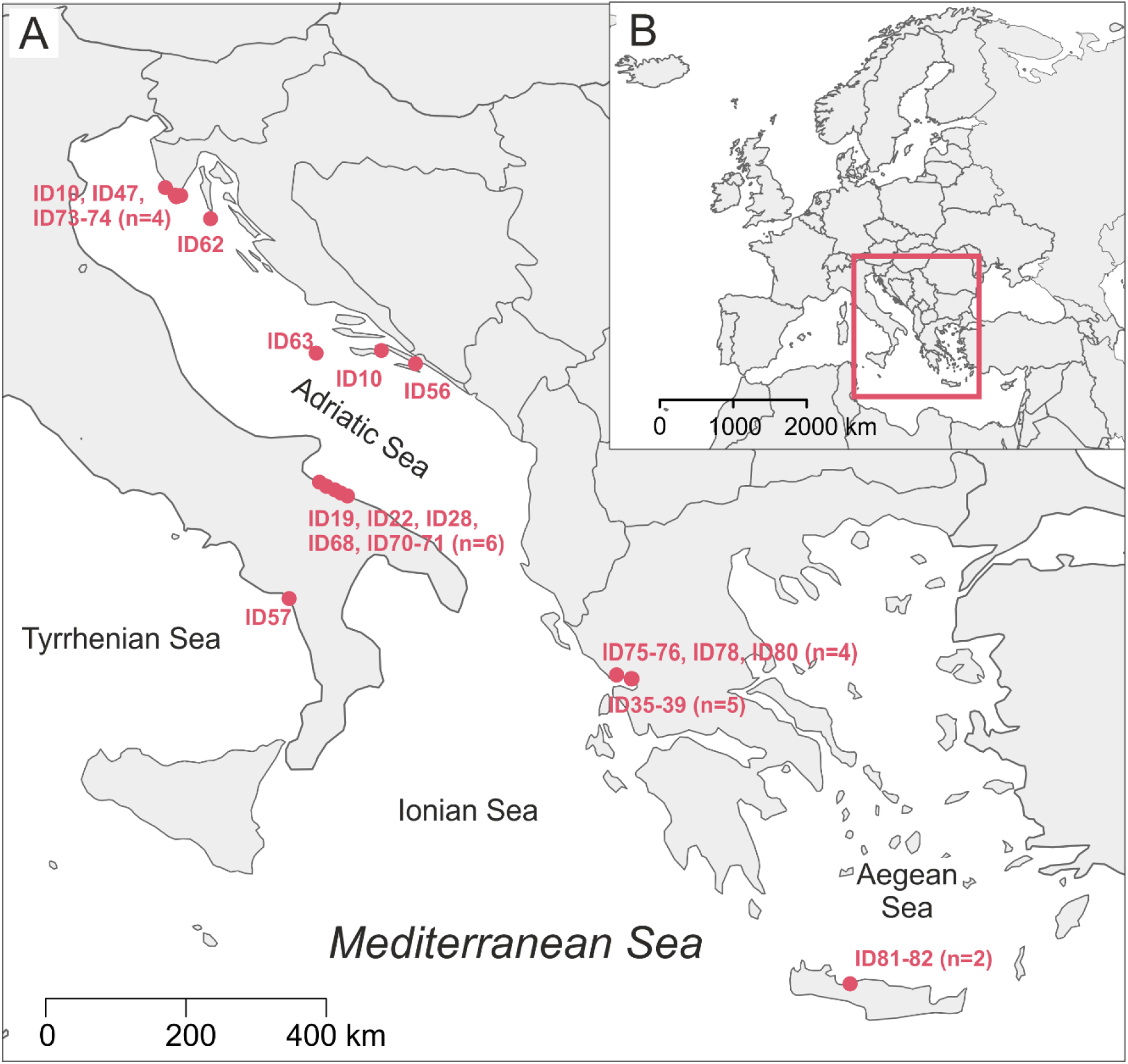
Map of the origin localities of sampled loggerhead turtles with indicated Turtle ID code (A); position of our study area in map of Europe (B).

**Table 1.** Turtle and sample information; n.a. indicates information not available; CCL – curved carapace length; asterisk (*) marks the straight carapace length (SCL) instead of curved carapace length (CCL).

| Turtle ID | Carapace sample ID | Skin sample ID | Origin sea | Origin locality | Sampling locality | Sampling date (DD.MM.YYYY.) | CCL (cm) | Turtle state |
| --- | --- | --- | --- | --- | --- | --- | --- | --- |
| ID010 | TB31 | TB32 | Adriatic Sea | CRO, Korčula | Pula Aquarium | 11.12.2018. | n.a. | Admitted |
|  | TB139 | TB140 | Adriatic Sea | CRO, Brijuni | Brijuni | 04.11.2019. | 69.7 | Wild |
| ID019 | TB49 | TB50 | Adriatic Sea | ITA, Barletta | Bari | 09.01.2019. | 50.7 | Wild |
| ID022 | TB55 | TB56 | Adriatic Sea | ITA, Barletta-Trani | Bari | 10.01.2019. | 72 | Wild |
| ID028 | TB73 | TB74 | Adriatic Sea | ITA, Barletta | Bari | 17.01.2019. | 74.5 | Wild |
| ID034 | TB89 | TB90 | Adriatic Sea | ITA, Bisceglie | Bari | 22.01.2019. | 72 | Wild |
| ID035 | GTB11 | GTB12 | Ionian Sea | GRE, Amvrakikos | Amvrakikos bay | 01.08.2018. | 78.6 | Wild |
| ID036 | GTB21 | GTB22 | Ionian Sea | GRE, Amvrakikos | Amvrakikos bay | 01.08.2018. | 51 | Wild |
| ID037 | GTB31 | GTB32 | Ionian Sea | GRE, Amvrakikos | Amvrakikos bay | 01.08.2018. | 69.6 | Wild |
| ID038 | GTB41 | GTB42 | Ionian Sea | GRE, Amvrakikos | Amvrakikos bay | 01.08.2018. | 58.5 | Wild |
| ID039 | GTB51 | GTB52 | Ionian Sea | GRE, Amvrakikos | Amvrakikos bay | 01.08.2018. | 53.2 | Wild |
| ID047 | TB115 | TB116 | Adriatic Sea | CRO, Kamenjak | Pula Aquarium | 08.05.2019. | 53.5* | Admitted |
| ID056 | TB117 | TB118 | Adriatic Sea | CRO, Ston | Pula Aquarium | 09.06.2019. | 74.0* | Admitted |
| ID057 | TB119 | TB120 | Tyrrhenian Sea | ITA, Maratea | Bari | 24.06.2019. | 77 | Admitted |
| ID062 | TB129 | TB130 | Adriatic Sea | CRO, Mali Lošinj | Lošinj | 30.07.2019. | 54 | Wild |
| ID063 | TB131 | TB132 | Adriatic Sea | CRO, Vis | Vis | 10.06.2019. | 24 | Wild |
| ID068 | TB145 | TB146 | Adriatic Sea | ITA, Molfetta | Bari | 25.07.2019. | 46.5 | Admitted |
| ID070 | TB149 | TB150 | Adriatic Sea | ITA, Molfetta | Bari | 24.07.2019. | 43 | Admitted |
| ID071 | TB151 | TB152 | Adriatic Sea | ITA, Margherita di Savoia | Bari | 23.10.2019. | 65.2 | Wild |
| ID073 | TB155 | TB156 | Adriatic Sea | CRO, Premantura | Pula Aquarium | 20.11.2019. | 32.2. | Admitted |
| ID074 | TB157 | TB158 | Adriatic Sea | CRO, Ližnjan | Pula Aquarium | 20.11.2019. | n.a. | Admitted |
| ID075 | GTB61 | GTB62 | Ionian Sea | GRE, Amvrakikos | Amvrakikos bay | 02.07.2019. | 64.5 | Wild |
| ID076 | GTB71 | GTB72 | Ionian Sea | GRE, Amvrakikos | Amvrakikos bay | 01.07.2019. | 62.9 | Wild |
| ID078 | GTB91 | GTB92 | Ionian Sea | GRE, Amvrakikos | Amvrakikos bay | 01.07.2019. | 55.3 | Wild |
| ID080 | GTB111 | GTB112 | Ionian Sea | GRE, Amvrakikos | Amvrakikos bay | 01.07.2019. | 66.5 | Wild |
| ID081 | GTB121 | GTB122 | Aegean Sea | GRE, Rethimno | Rethimno bay | 10.07.2019. | n.a. | Wild |
| ID082 | GTB131 | GTB132 | Aegean Sea | GRE, Rethimno | Rethimno bay | 14.07.2019. | n.a. | Wild |

### 2.2 DNA analysis

The DNA isolation and sequencing were performed twice, in 2019 (20 samples from ID10, ID19-39) and in 2020 (34 samples from ID10, ID47-82). The DNA was extracted from 0.25 g of an ethanol-free sample in duplicates using the DNeasy PowerSoil kit (QIAGEN, RRID:SCR_008539). The extraction protocol followed the manufacturer’s guidelines with several modifications (as described below). Samples were transferred into the PowerBead tubes and incubated in a sonicator at 50 °C at 35 kHz for 15 minutes. The incubation times for C1, C2, and C3 solutions were extended (30 minutes at 65 °C for C1 and 15 minutes at 4 °C), and bead-beating was replaced with horizontal vortexing on IKA VXR basic Vibrax shaker (10 minutes at maximum speed of 2200 rpm). The DNA was eluted with 50 μl of DNase-free molecular grade water (incubated at room temperature for 2 minutes). The quantity and purity of extracted DNA were measured by NanoDrop ND-1000 V3.8 spectrophotometer (ThermoFisher). The extracted DNA samples were sent for 2×250 bp paired-end sequencing (Illumina MiSeq System, RRID:SCR_016379) of the 16S rRNA gene V4 region by 515F (5′-GTGYCAGCMGCCGCGGTAA-3′) and 806R (5′-GGACTACNVGGGTWTCTAAT-3′) primers (Apprill et al., 2015; Parada et al., 2016), and the 18S rRNA gene V4 region by eukV4F (5′-CCAGCASCYGCGGTAATTCC-3′) and zigeukV4R (5′-ACTTTCGTTCTTGATYRATGA-3′) primers (Stoeck et al., 2010; Piredda et al., 2017) at Molecular Research MrDNA (Shallowater, TX, USA).

### 2.3 Sequence data processing and analysis

Sequences obtained from MrDNA were processed by FASTqProcessor (MrDNA), and all non-biological sequences were removed prior to exporting the data in QIIME2-readable format (“EMP protocol” multiplexed paired-end fastq format). The sequences were then imported to the QIIME2 (RRID:SCR_021258) environment, versions 2020.6 for 16S and 2021.4 for 18S (Bolyen et al., 2019). Demultiplexing of sequences was done by q2-demux plugin. DADA2 (q2-dada2 plugin) was used for sequence denoising (Callahan et al., 2016). 18S rRNA sequences were truncated at 220 bp for forward and reverse sequences. Sequence alignment was performed with MAFFT (Katoh et al., 2002) and a phylogenetic tree was constructed with fasttree2 using q2-phylogeny (Price et al., 2010). Taxonomy was assigned to amplicon sequence variants (ASVs) via q2-feature-classifier (Bokulich et al., 2018) classify-sklearn naïve Bayes taxonomy classifier against the SILVA v.138 (99% 505F-806R nb classifier) (Quast et al., 2013) and PR2 4.13.0 (Guillou et al., 2013; Campo et al., 2018) databases for 16S and 18S datasets, respectively. Prior to downstream analyses, mitochondria and chloroplast sequences were filtered from the 16S dataset, and metazoan and macroalgal sequences were filtered from the 18S dataset. For alpha and beta diversity analyses, we rarefied the 16S dataset to the sampling depth of 34 000 and the 18S dataset to 10 000 based on rarefaction curves (q2-diversity plugin). We calculated two alpha diversity indices via q2-diversity: observed ASVs (features) and Faith’s phylogenetic diversity (PD) index (Faith, 1992) for both 16S and 18S datasets, and made visualizations using boxplots. Beta diversity was estimated using three distance matrices via q2-diversity: Bray-Curtis, weighted UniFrac (Lozupone et al., 2007) and robust Aitchison’s distance (Aitchison and Shen, 1980; Aitchison and Ho, 1989; Martino et al., 2019). Principal Coordinates Analysis (PCoA) plots for Bray-Curtis and weighted UniFrac distance were produced using the q2-diversity plugin, and Robust Aitchison Principal Components Analysis (rPCA) was performed on non-rarefied data via DEICODE plugin (Martino et al., 2019). To compare the 16S and 18S datasets, we plotted the first principal coordinate (PC1) of each dataset’s robust Aitchison’s distance and performed the Procrustes analysis via q2-diversity. The permutational multivariate analysis of variance (PERMANOVA) with the q2-diversity plugin was used to test for significant differences between sample groups. Turtles from the Aegean and Tyrrhenian Seas were excluded from PERMANOVA calculations for “Origin Sea” due to the low number of samples in these two groups. For PERMANOVA statistic, turtles from Adriatic Sea were divided on East Adriatic (Croatian samples) and West Adriatic (Italian samples). PERMANOVA tested factor “Season” was obtained as following: samples obtained in spring and summer are put into “warm” category, while samples from autumn and winter are put into “cold” category. Data visualizations were made using ggplot2 (RRID:SCR_014601) (Wickham, 2016), phyloseq (RRID:SCR_013080) (McMurdie and Holmes, 2013), vegan (RRID:SCR_011950) (Oksanen et al., 2020), and pheatmap (RRID:SCR_016418) (Kolde, 2019) within R Studio (R Project for Statistical Computing, RRID:SCR_001905). The relative abundance of different groups of samples was calculated as a sum of the ASV count of selected taxon and then divided by the total sequence number in that sample group. Community composition was summarized by heatmaps (Fig. 2) produced based on centred log-ratio (clr) transformed data from ASV counts.

**Figure 2.**
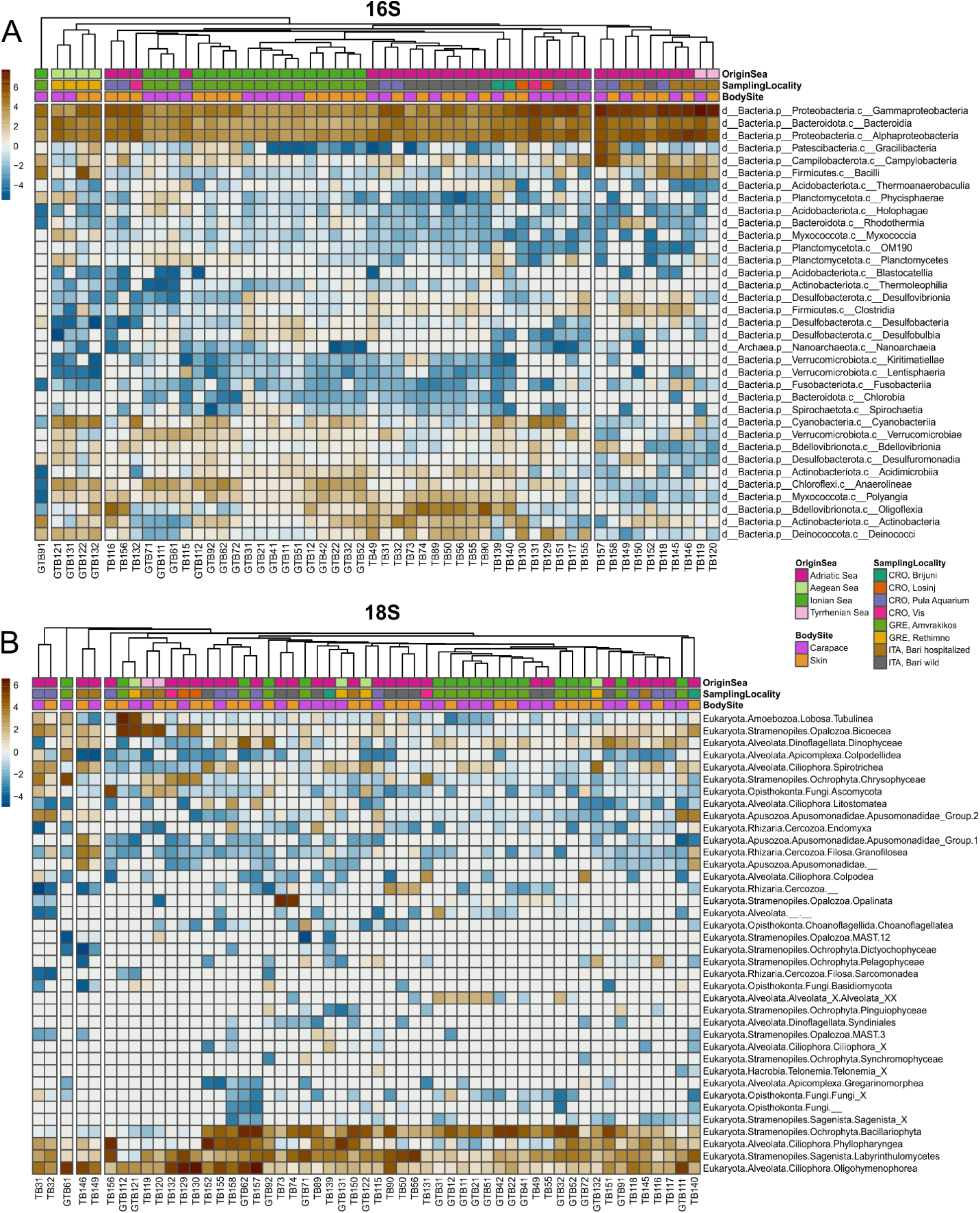
Heatmaps of surface microbiome of loggerhead sea turtles; showed taxa are more than 1% abundant in at least one sample.

## 3 Results

High throughput sequencing of 54 samples yielded 6,242,910 high quality 16S and 1,675,191 18S sequences. Median frequency per sample was 102,920.0 (min. 34,669.0; max. 257,399.0) for 16S while median frequency per sample for 18S was 20,048.5 (min. 1,634.0; max. 123,842.0). Sequences obtained for 16S and 18S were denoised to 17,636 ASVs and 1,917 ASVs, respectively (Tables S1, S2 and S3).

### 3.1 Community composition

#### 3.1.1 Prokaryotic microbiota

The prokaryotic community showed the dominance of bacterial over archaeal taxa. The most abundant bacterial phylum in all samples was Proteobacteria, followed by Bacteriodota, Bdellovibrionota and Cyanobacteria (Figs 2 and 3). Classes Gammaproteobacteria, Alphaproteobacteria and Bacteroidia dominated in all samples (Fig. 2). Family *Rhodobacteraceae* was more abundant overall than any other prokaryotic family, followed by *Moraxellaceae* and *Pseudoalteromonadaceae* (Table S4).

**Figure 3.**
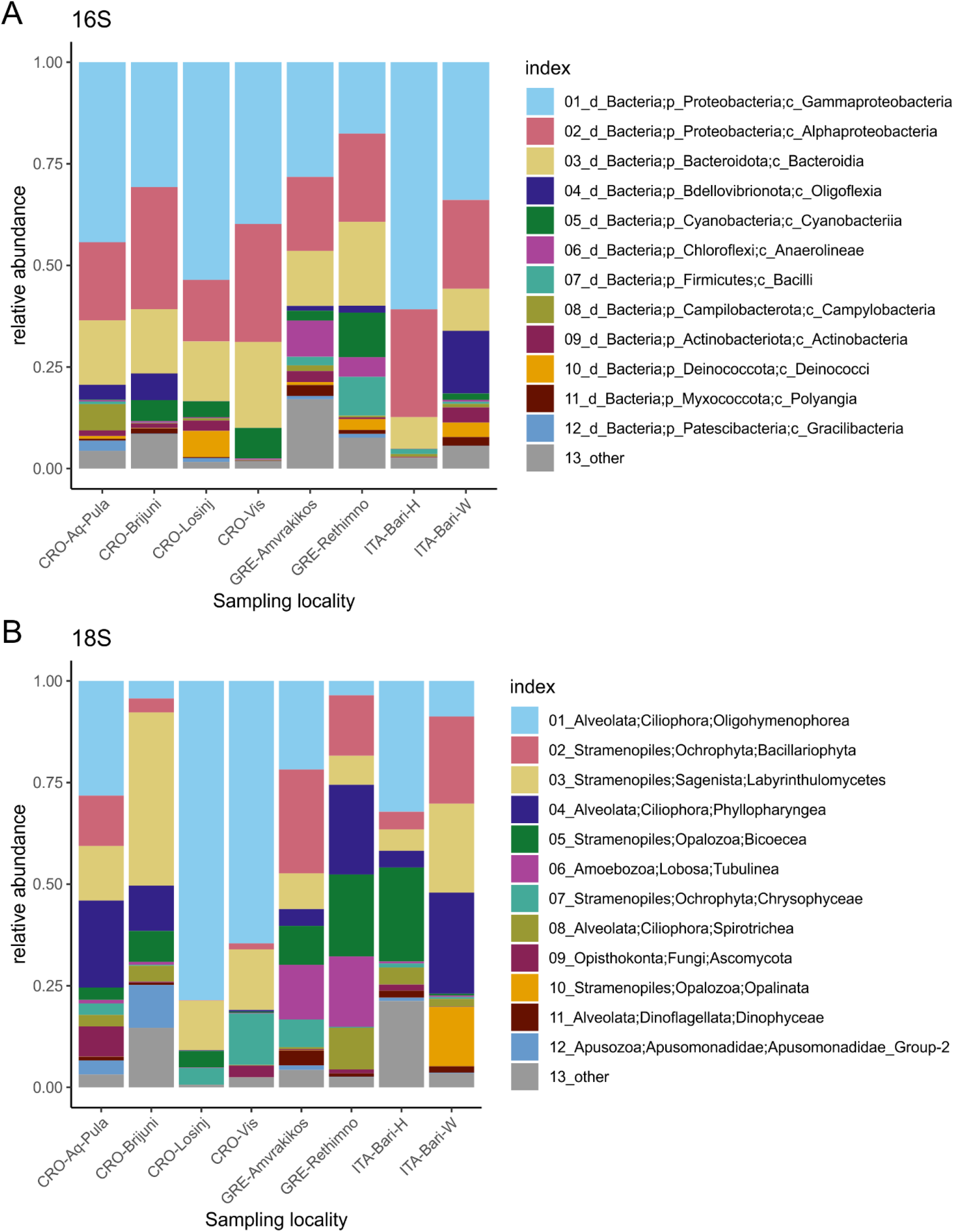
Relative abundances of 12 most abundant microbial taxa present on loggerhead sea turtles from the same sampling locality 16S dataset (up), 18S dataset (bottom).

There were five core features (ASVs) identified in 100% of samples in the 16S dataset, four belonging to class Gammaproteobacteria and one to Oligoflexia (Table S5). These features are classified as an uncultured bacterium from order Oligoflexales, genus *Pseudoalteromonas*, an unidentified ASV from class Gammaproteobacteria, an uncultured bacterium from family *Sedimenticolaceae*, and an uncultured bacterium from family *Saccharospirillaceae*. In carapace samples, additional five core features were identified in 100% of the samples classified as genus *Vibrio* (Gammaproteobacteria), BD1-7 clade (family *Spongiibacteraceae*, Gammaproteobacteria), an uncultured bacterium from family *Arcobacteraceae* (Campylobacteria), genus *Deinococcus* (Deinococci), and genus *Halarcobacter* (Campylobacteria). In skin samples, additional seven core features were identified and classified as an uncultured bacterium from family *Nannocystaceae* (Polyangia, Myxococcota), family *Rhodobacteraceae* (Alphaproteobacteria), an uncultured bacterium from genus *Psychrobacter* (Gammaproteobacteria), an uncultured bacterium from genus *Ahniella* (Gammaproteobacteria), genus *Tenacibaculum* (Bacteroidia), family *Stappiaceae* (Alphaproteobacteria), and genus *Poseidonibacter* (Campylobacteria).

#### 3.1.2 Eukaryotic microbiota

The most abundant supergroups of micro-eukaryotes in the dataset were Alveolata and Stramenopiles. The dominant classes included Oligohymenophorea, Bacillariophyta, Labyrinthulomycetes and Phyllopharyngea. The class Opalinata was highly prevalent in samples TB73 and TB74 (turtle ID28). Samples TB119 (carapace) and TB120 (skin) (from turtle ID57, the only animal sampled in the Tyrrhenian Sea) were dominated by Biocoeca. A high abundance of Ascomycota (Fungi) was recorded in the skin sample TB156 (Adriatic Sea), while *Chrysophyceae* were particularly abundant in the carapace sample GTB61 (Ionian Sea).

One core feature, belonging to the genus *Zoothamnium* (Oligohymenophorea, Ciliophora), was identified in all samples within the 18S dataset. An additional core feature of the carapace samples was found to be *Nitzschia communis* (Bacillariophyta). No additional core features were shared by all skin samples (Table S5).

At the genus level (level 7 in the PR2 database), apart from the above-mentioned *Zoothamnium*, a taxon assigned to the level of “Raphid-pennate” group (Bacillariophyta) was found in all biofilm samples. In all carapace samples, *Nitzschia* (Bacillariophyta) and *Labyrinthula* (Labyrinthulomycetes) were identified as additional core genera. In 95% of all biofilm samples, the following core features were identified at the genus level: “Raphid-pennate” group (Bacillariophyta), *Zoothamnium* (Oligohymenophorea, Ciliophora), *Nitzschia* (Bacillariophyta) and *Labyrinthula* (Labyrinthulomycetes). In 95% of carapace samples, an additional core genus, *Caecitellus* (Opalozoa), was found. Four additional core genera were detected in 95% of skin samples: *Thraustochytrium* (Labyrinthulomycetes), *Uronema* (Oligohymenophorea), *Labyrinthulaceae* X (Labyrinthulomycetes), and *Fistulifera* (Bacillariophyta).

#### 3.1.3 Prokaryotic photoautotrophs

Within the 16S dataset, Cyanobacteria were the most abundant photoautotrophic prokaryotes, with *Phormidesmiaceae* and *Paraspirulinaceae* being the most abundant (Fig. 4). On average, Cyanobacteria comprised 3% of all ASV sequences. In individual samples, this group accounted for 0.01-19.77% of all sequences (Fig. 4B). *Phormidesmiaceae* and *Xenococcaceae* dominated in carapace samples, whereas *Paraspirulinaceae* and *Phormidesmiaceae* were most abundant in skin samples. Many of the detected cyanobacterial sequences remained unclassified (Fig. 4A, pink bars).

**Figure 4.**
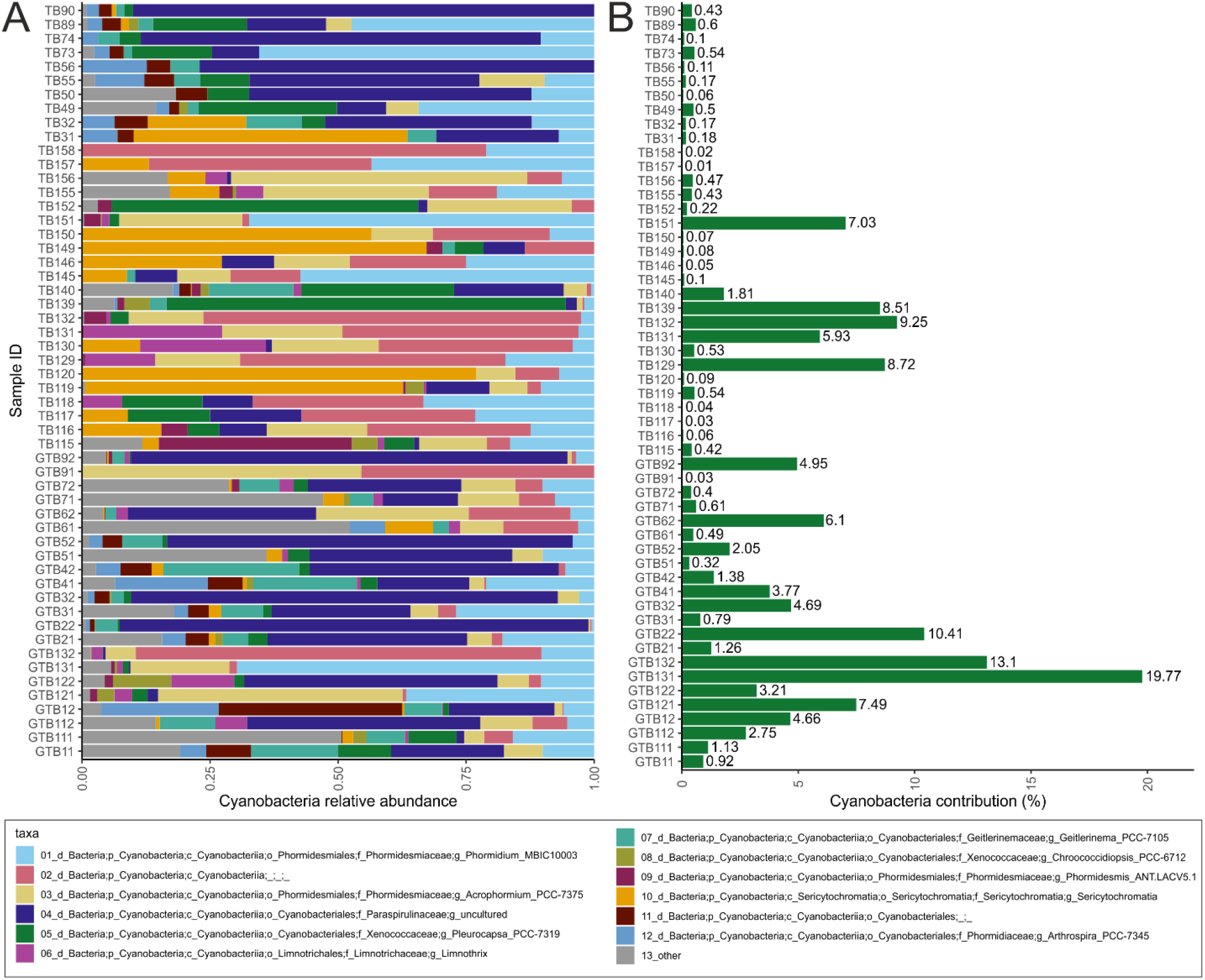
Relative abundances within the cyanobacterial group (A); contribution of Cyanobacteria to total prokaryotic community within a sample based on relative abundance (B).

### 3.2 Alpha diversity

Alpha diversity indices for community richness (observed ASVs) and diversity (Faith’s Phylogenetic Diversity (PD) index) were highly variable and ranged from 127 to 2833 (richness) and from 12.68 to 135.65 (diversity) for the prokaryotic community (Fig. 5A and B). Prokaryotic communities from turtles in different seas and from different body sites (“Origin Sea” and “Body Site” categories as shown in Table 1) showed significant differences (Kruskal-Wallis H test, p>0.05). Within the 16S dataset, carapace samples showed higher median values of richness (1032) and diversity (52.23) than skin samples (richness 811; diversity 46.47). The highest median values of Faith’s PD were observed for samples from the Ionian Sea (75.36), followed by the Aegean (59.39) and Adriatic Seas (42.17). The lowest Faith’s PD values were recorded for the hospitalised turtle from the Tyrrhenian Sea (20.69; ID057). Microbial eukaryotes’ community ASVs richness ranged from 44 to 197, and diversity values ranged from 9.47 to 25.53 (Fig. 5C and D) which is considerably lower comparing to the prokaryotes. The highest median value of micro-eukaryotic ASV richness was observed for the Adriatic Sea (124.5), followed by Ionian Sea (112), Aegean Sea (89.5), and Tyrrhenian Sea (82). The highest median value of micro-eukaryotic Faith’s PD was observed for Ionian Sea (17.11), followed by Adriatic Sea (16.91), Tyrrhenian Sea (15.76) and Aegean Sea (15.35), similar to the prokaryotic communities. Carapace microbial eukaryotes showed higher median values of ASVs richness (118) and diversity (17.13) than skin samples (richness 93, diversity 14.75); however, no significant differences between the different seas or body sites were observed.

**Figure 5.**
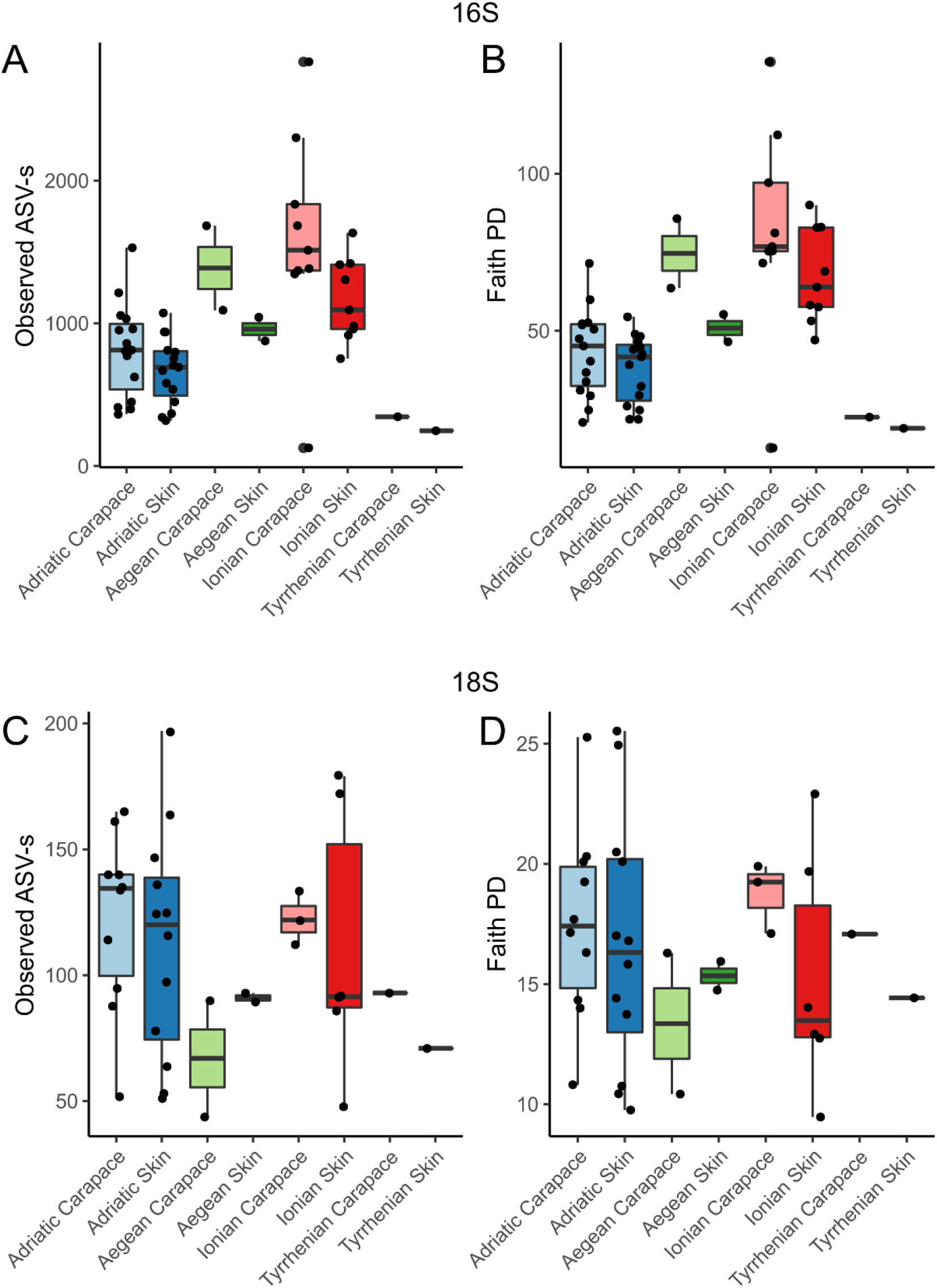
Observed ASV richness and Faith’s phylogenetic diversity index for 16S and 18S datasets for skin and carapace microbiome of loggerhead sea turtles sampled at 4 locations in the Mediterranean.

### 3.3 Beta diversity

Principal Components Analyses of robust Aitchison distance (rPCA) indicate groupings based on sampling locality and body site for prokaryotes (Figure 6A) and eukaryotes (Figure 6B). For the 16S dataset (Figure 6A) we can observe groupings based on sampled body site (carapace on the right and skin on the left) and sampling locality. ASVs that drive those groupings belong to uncultured Oligoflexales, Nannocystales, *Rhodobacteraceae, Saccharospirillaceae, Pseudoalteromonas* and *Vibrio*. For the 18S dataset (Fig 6B) we cannot observe clear groupings based on body site but there is an indication of samples grouping based on sampling locality. ASVs that drive the sample distribution for micro-eukaryotic communities (Fig. 6B) belong to Ciliophora (*Zoothamnium* sp., Sessilida, *Uronema marinum, Uronema nigricans, Ephelota gigantea, Aspidiscida steini*), *Nitzschia communis* (Bacillariophyta) and *Cafeteria roenbergensis* (Bicoecea). To gain insight into the whole epi-microbiome (prokaryotic and eukaryotic) we combined the principal components (PC1s) of the rPCA for 16S and 18S datasets where clear groupings based on sampling locality can be distinguished (Fig. 7).

**Figure 6.**
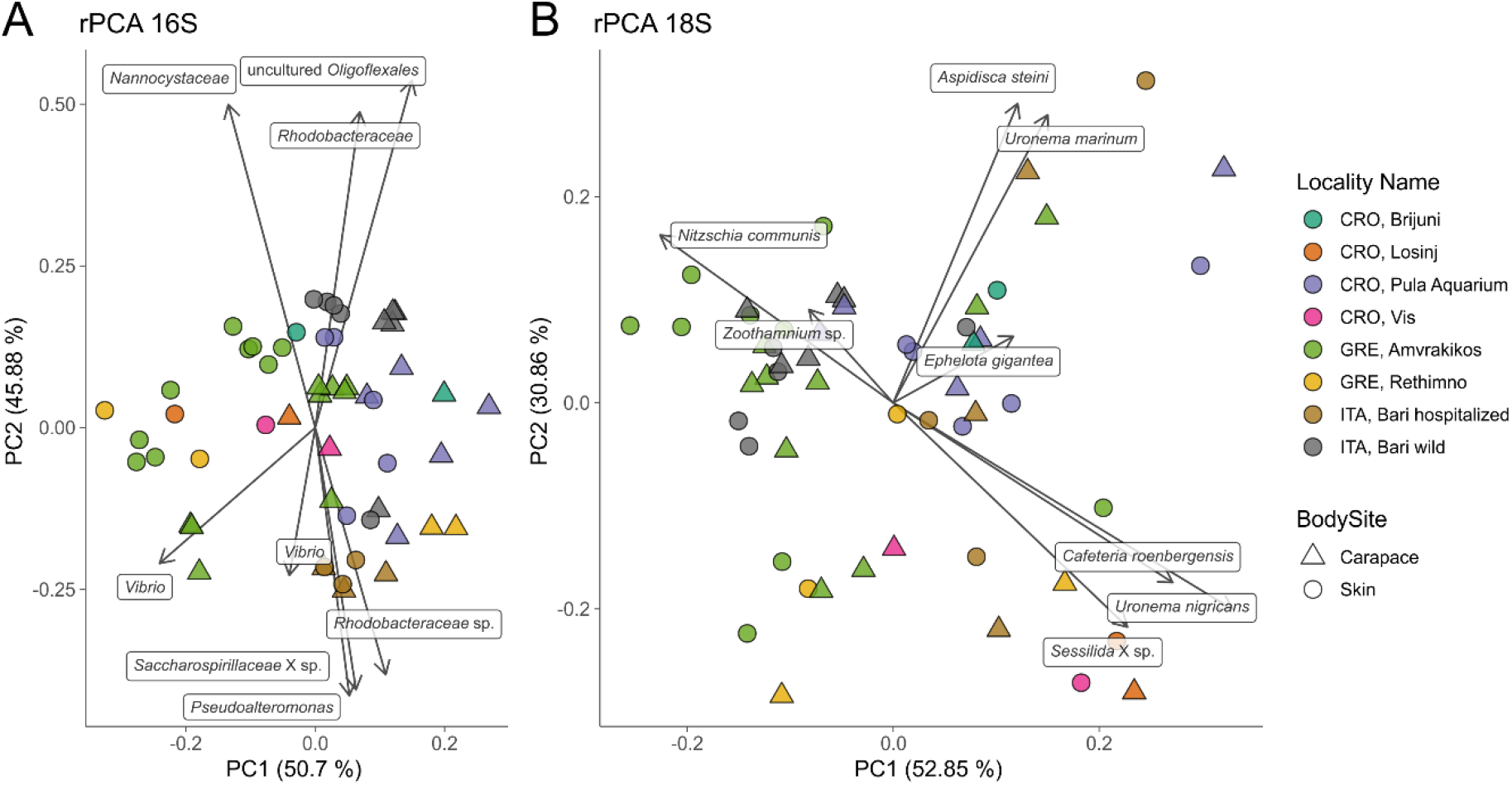
Principal component analysis (PCA) biplot of robust Aitchison distance for prokaryotic (16S, left) and eukaryotic diversity (18S, right); arrows indicate individual highly ranked ASVs that contribute to the displayed positions of the samples; lowest taxonomic assignment of each ASV is written in textboxes at the end of each arrow.

**Figure 7.**
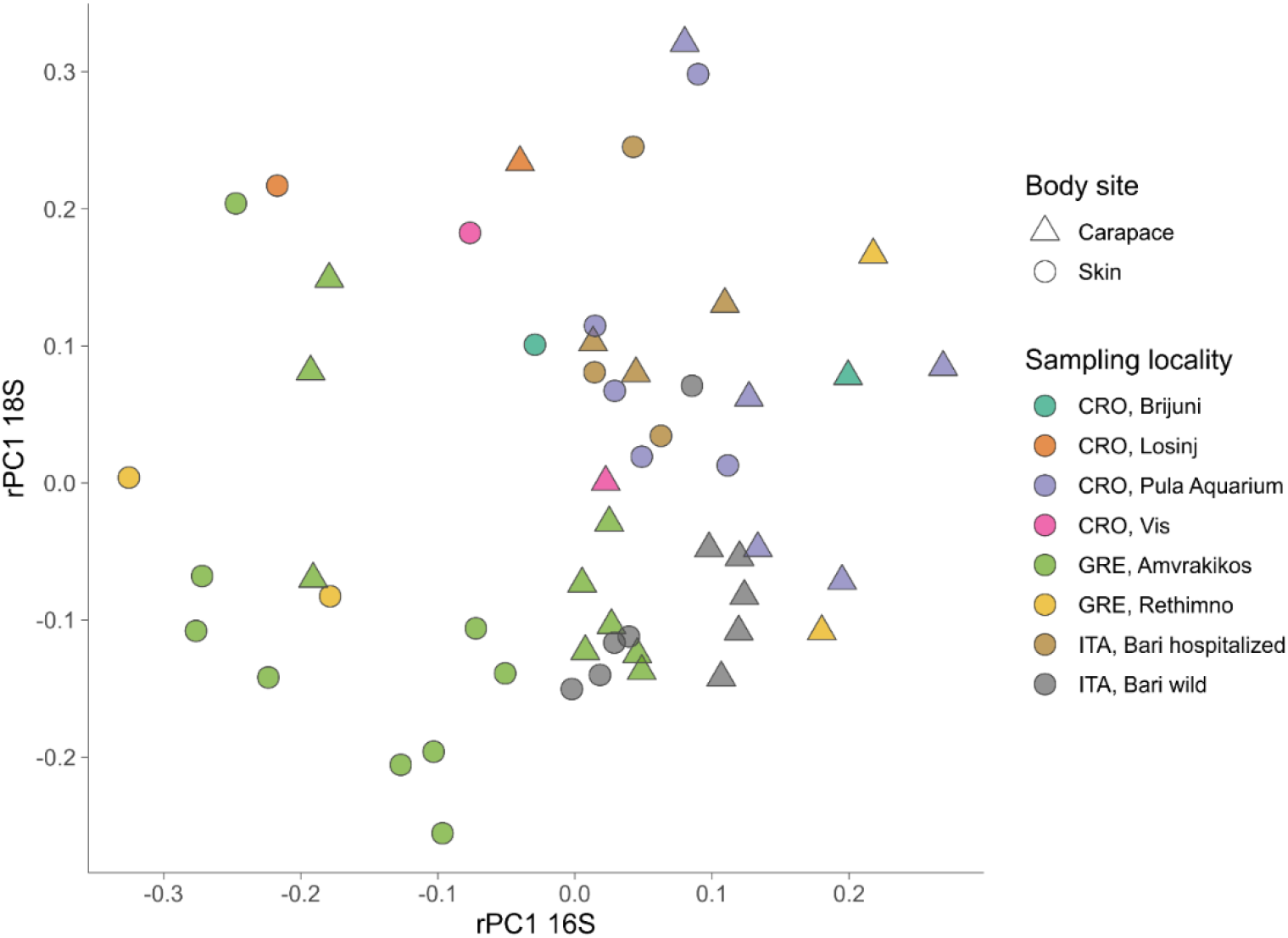
Combination plot of the first principal component (PC1) of PCAs based on robust Aitchison distance matrix for 16S (x-axis) and 18S (y-axis). Sampling locality are indicated by color, body sites are indicated by shape.

To compare and detect any congruence between the prokaryotic and eukaryotic communities the rPCA ordinates of both datasets were compared by the Procrustes analysis (Figure S2) which showed the prokaryotic and eukaryotic dataset congruence is low (m2 = 0.93095, p = 0.043).

PERMANOVA results (Table 2) show that there is a significant difference (p < 0.05) between the prokaryotic communities of skin and carapace body sites, the seas of origin (only Adriatic and Ionian), turtle state (wild vs. admitted), and sampling season (warm vs. cold). The only non-significant value was detected between body site groups for unweighted UniFrac. The highest pseudo-F values for all distance matrices were observed between “Origin Sea” categories.

**Table 2.**
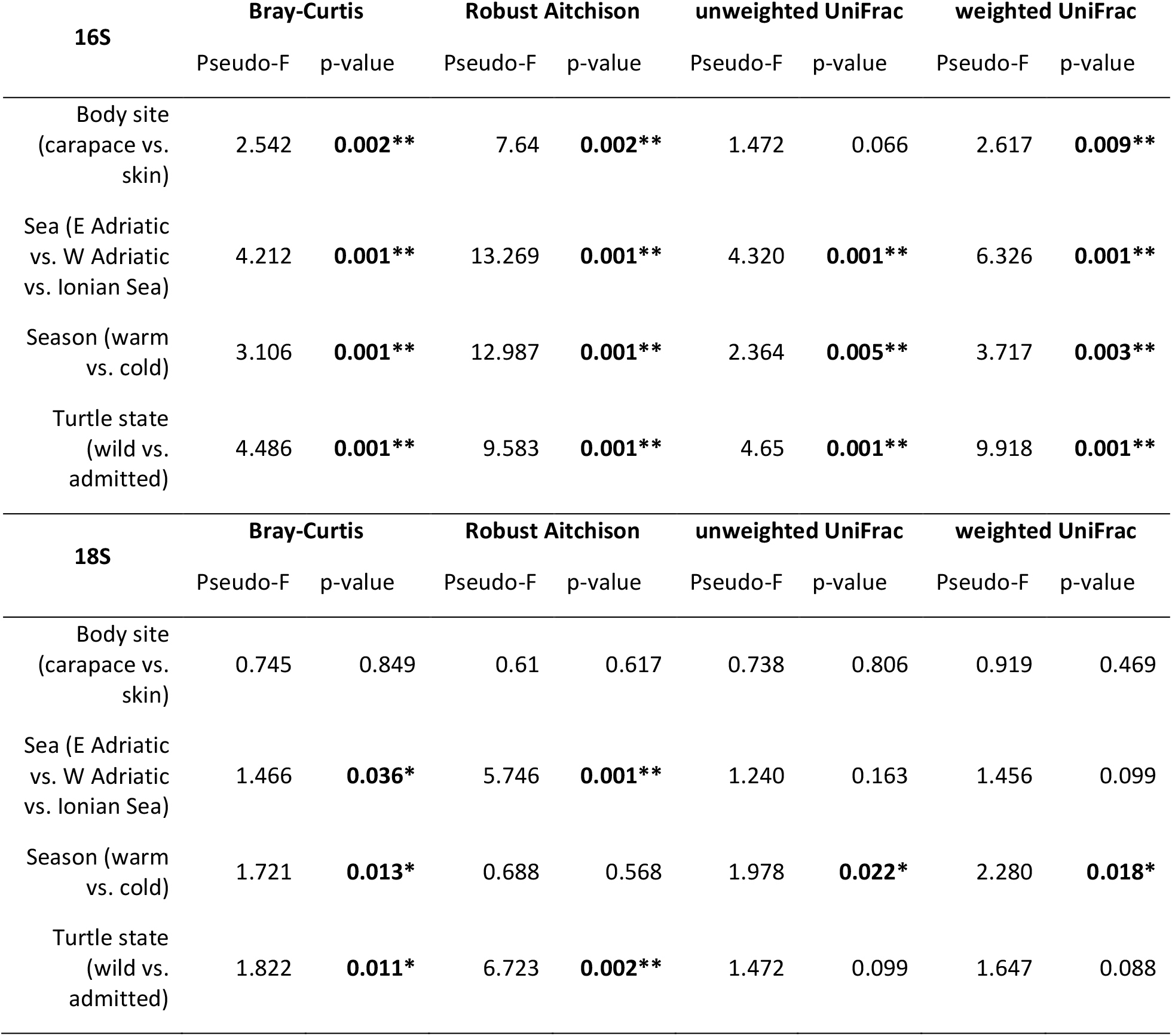
Permutational multivariate analysis of variance (PERMANOVA) for Bray-Curtis, Robust Aitchison, unweighted and weighted UniFrac distance metrics. Significance levels are indicated by an asterisk: p ≤ 0.05*, p ≤ 0.01** with all significant values bolded.

PERMANOVA on eukaryotic communities showed no significant differences between sampled body sites. Significant differences between origin seas and turtle states were observed for Robust Aitchison and Bray-Curtis distances. According to all but one distance metrics tested, PERMANOVA showed a significant difference between sampling seasons (Table 2).

## 4 Discussion

In this study we provide insights into the epi-microbiota of loggerhead sea turtles using a combined 16S and 18S metabarcoding approach. Our results show that overall prokaryotic microbiota is dominated by a few classes of bacteria (Gammaproteobacteria, Alphaproteobacteria and Bacteroidia) and that the communities may differ depending on multiple extrinsic and intrinsic factors, which has been previously described in studies on other aquatic animals (as reviewed in Apprill, 2017). On the other hand, in spite of eukaryotic microbiota showing high heterogeneity, core taxa such as Oligohymenophorea, Bacillariophyta, Labyrinthulomycetes, and Phyllopharyngea were commonly present in the majority of samples. Despite the sampled turtles coming from different locations in the Mediterranean Sea, varying in age and health conditions, and being sampled in different seasons forming a diverse dataset, it is clear that several of the tested factors influenced their surface microbial community composition. Prokaryotic communities seem to be affected by the locality of origin, body site, turtle state, and sampling season while the eukaryotic microbiota followed a similar pattern, although to a lesser extent, and without detected differences between body sites.

The highest microbial diversity was observed on Ionian turtles from the lagoonal complex of the Amvrakikos Gulf, that is one of the most important and productive lagoonal complexes in Greece. The lagoonal shallow coastal aquatic systems, with a maximum depth of 65 m, are separated from the sea by sediment barriers and connected to it through channels, often characterized by salinity fluctuations and development of low dissolved oxygen conditions. While the Amvrakikos Gulf offers a rich neritic foraging ground for subadult and adult loggerheads (Rees et al., 2013), the second locality with highest diversity is the Rethimno bay in northern Crete (Greece) that is an important nesting site for adult females (Margaritoulis and Rees, 2011). Low diversity and richness of turtle-associated microbial communities from the Tyrrhenian Sea could be due to short continental shelf and lower availability of the rich benthic environment as a possible source of microbes which could colonize the loggerhead’s body.

Prokaryotic diversity and richness of carapace samples was consistently higher than those of skin samples which could be explained by the large and rigid surface of the carapace covered by keratinous scutes that could allow for easier attachment and colonization of diverse microbes. Compared to the carapace, the skin of the neck and flippers (sampled in this study) is prone to higher mechanical disturbance caused by the turtle’s movements. Parks et al. (2020) reported a higher diversity and richness of microbial communities on the freshwater turtles’ carapace in comparison to the plastron, and provide the movement of the turtles as one of the possible explanations. Furthermore, Blasi et al., (2022) reported significant differences between microbial communities of differently positioned carapace scutes. The difference in bacterial community composition of anterior and posterior scutes of the sea turtle carapace might have been caused by different abiotic (hydrodynamics or sun exposure) and biotic factors (uneven distribution of macroorganisms across the carapace) affecting those areas (Blasi et al., 2022). The epi-microbiota of three juvenile loggerheads from the Tyrhhenian Sea harboured Firmicutes and Proteobacteria as the most prevalent phyla (Blasi et al., 2022). Contrastingly, in our dataset Proteobacteria were found to be the most abundant while Firmicutes were not among the highly abundant phyla. The most abundant bacterial family was *Rhodobacteraceae* which is known to be widely distributed in marine benthic habitats (Pohlner et al., 2019). Although the metabolic variety within *Rhodobacteraceae* is great, they are mainly aerobic photo- and chemoheterotrophs, and purple non-sulfur bacteria that are known for anaerobic photosynthesis (Pujalte et al., 2014). Interestingly, we observed uncultured members of *Psychrobacter* and *Tenacibaculum* genera on all of the skin samples which were also reported as a part of the core microbiome on the humpback whales (Bierlich et al., 2018). This raises a question about *Psychrobacter* and *Tenacibaculum* genera members’ dependence on animal skin metabolites, possibly making them mutualistic or commensal to marine animals. It is worth mentioning that some species of *Tenacibaculum* are known as pathogens on fish skin (Nowlan et al., 2020), however we cannot be certain of the exact ecological role on the turtle skin without further research.

The micro-eukaryotic taxa which were dominant in most samples, ciliates Oligohymenophorea and Phyllopharyngea (Alveolata), are mainly free-living heterotrophs that could possibly graze on the other microbes colonizing the turtles’ surfaces. Commonly found Stramenopiles were mostly represented by diatoms (Bacillariophyta) and Labyrinthulomycetes (marine fungus-like organisms that produce filamentous webs for nutrient absorption). Diatoms are photosynthesizing microalgae with characteristic silica shells (Round et al., 1990) known for being among the first colonizers of submerged surfaces including marine vertebrates (Hooper et al., 2019) Our results show that diatoms are one of the major eukaryotic groups present on sea turtles’ bodies and the most dominant phototrophs in those communities. Contrary to prokaryotic community, differences between carapace and skin community were not detected for micro-eukaryotes. This is also not in congruence with the morphological study on diatoms from loggerheads of where they reported higher diversity and richness of carapace than in skin diatom community. Common epiphytic and epipelic diatom genera were found abundant on carapace while putatively epizoic taxa were dominating in skin diatom samples (Van de Vijver et al., 2020).

Light availability on the sea turtle surfaces enables the development of phototrophic microbes that cannot be found as a part of the endozoic microbiome. Moreover, unlike endozoic microbial communities which are dependent on nutrient inflow from the host, epizoic communities are probably dependent mostly on the nutrients available in the surrounding environment and from the primary producers in those communities. Previous morphology-based studies on diatoms revealed high abundance, diversity, and uniqueness of diatom species associated with the sea turtles (Majewska et al., 2015; Rivera et al., 2018; Van de Vijver et al., 2020). Besides diatoms, photoautotrophic *Chrysophyceae* and *Dinophyceae* were detected in noticeable abundances, and Chrysophyte stomatocysts of unknown species were previously reported on the sea turtle carapace (Pang et al., 2021). Additionally, Labyrinthulomycetes that were the third most abundant taxon in our samples are known to be mainly decomposers or, rarely, parasitic (Tsui et al., 2009) with recently emphasized importance in carbon sequestration (Bai et al., 2021). Either turtle- or microbe-derived particulate carbon (photoautotrophs or heterotrophs) could provide Labyrinthulomycetes with significant amounts of energy sources leading to their high relative abundance across samples.

Prokaryotic photoautotrophic communities are dominated by Cyanobacteria with the most common in our samples being filamentous cyanobacteria like *Phormidium* and *Leptolyngbya* (*Acrophorium*), both known for cyanotoxin production (Frazão et al., 2010; McAllister et al., 2016; Li et al., 2019). Due to climate change and rising temperatures, toxic cyanobacterial blooms are predicted to become a progressively serious problem (O’Neil et al., 2012) and it has been observed that cyanobacterial toxic compounds can interfere with composition and function of animal intestinal microbiome (Duperron et al., 2019; Li et al., 2019; Sehnal et al., 2021). Other common genera we found include *Pleurocapsa, Limnothrix, Geitlerinema, Chroococcidiopsis*, and *Phormidesmis*. Blasi et al. (2022) also highlighted the presence of Cyanobacteria on anterior scutes, specifically families *Pseudanabenaceae* and *Rivulariaceae* while *Phormidesmiaceae, Paraspirulinaceae* and *Xenococcaceae* were prevalent in our dataset. All reported cyanobacterial genera in our study are commonly found in marine benthic habitats forming colonies and cyanobacterial mats (Komárek et al., 2014). Sea turtles seem to provide additional surfaces for cyanobacterial colonization and could act as a highly mobile reservoir with unknown implications for the host’s health and effects on the environment.

It should be noted, however, that observed significant differences in multiple groupings of microbial communities in this study could be explained by overlapping metadata categories (e.g., wild animals being sampled mostly in Greece and during summer months) that could not be controlled for within our study design due to the unpredictability/stochasticity of opportunistic sampling. Additionally, reference databases play an important role in investigating microbial eukaryotes, as we cannot grasp the full diversity of micro-eukaryotes through metabarcoding alone because of a lack of sequenced representatives and eukaryotes often being overlooked as a part of microbial communities (Lind and Pollard, 2021 and references therein). In our study, a major portion of the cyanobacterial ASVs could not be properly identified via metabarcoding as the current version of SILVA reference database taxonomy is based on Bergey’s Manual of Systematic Bacteriology (Boone et al., 2001) in which cyanobacterial taxonomy higher than genus is not defined. Therefore, SILVA and Genome Taxonomy Database (GTDB) (Quast et al., 2013) proposed their own names for some taxa based on 16S rRNA phylogeny that is not in agreement with the currently valid cyanobacterial taxonomy in the CyanoDB database (Komárek et al., 2014). As microbial eukaryotes and cyanobacteria are an important part of microbial communities associated with Mediterranean loggerhead sea turtles, further efforts in their characterization are needed to reconcile multiple taxonomy databases and better understand the turtle-associated taxa and their possible effects on the host

The Mediterranean loggerheads are widely distributed large hard-shelled top predators, and a highly migratory species which occupies different marine habitats at different life stages. Their major ecological role in bioturbation, energy flow, trophic status, mineral cycling, soil dynamics and connectivity between habitats makes it a keystone species in Mediterranean marine environment (Casale et al., 2018). This research brings us one step closer to much needed understanding of the complexity of microbial communities associated with loggerheads and wild animals in general. We show in this study that microbial communities of loggerhead sea turtles are rich and highly diverse with reservoirs of microbial taxa potentially important both for turtles’ and the ecosystem’s state. Moreover, DNA-based surveys focusing on epizoic prokaryotic and eukaryotic microbiota could prove to be a valuable addition to non-invasive methods for monitoring the status of endangered marine species and their environment.

## 5 Acknowledgements

We are thankful to Milena Mičić, Karin Gobić Medica and the rest of the staff from the Marine Turtle Rescue Center (Aquarium Pula) as well as Mateja Zekan and Draško Holcer from Blue World Institute for sample collection and to Hrvoje Višić for help with laboratory work.

## 6 Data Availability Statement

The amplicon sequence data is deposited in the European Nucleotide Archive (ENA) under accession numbers PRJEB51458 for 16S and PRJEB51472 for 18S.

## 7 Ethics Statement

Sampling was performed in accordance with the 1975 Declaration of Helsinki, as revised in 2013 and the applicable national laws. The sampling at the Sea Turtle Clinic (Bari, Italy) was conducted with the permission of the Department of Veterinary Medicine Animal Ethic Committee (Authorization # 4/19), while sampling in Croatia was done in accordance with the authorization of the Marine Turtle Rescue Centre by the Ministry of Environment and Energy of the Republic of Croatia. Sampling activities in Greece were carried out with permission from the Hellenic Ministry of Agriculture and Environment.

## 8 Conflict of interest

The authors declare that the research was conducted in the absence of any commercial or financial relationships that could be construed as a potential conflict of interest.

## 9 Author Contributions

SB, RG, and RM designed the study; LK, KF, AT, MC, AB, AP and SB collected the samples; KF and MM carried out the laboratory work; LK and KF conducted the bioinformatics, statistical analyses, and data visualization and interpretation; LK wrote the first draft of the manuscript; SB conceived project and obtained funding; all authors revised the paper and approved the final version of the manuscript.

## 10 Funding

The study has been fully supported by the Croatian Science Foundation under the project number UIP-2017-05-5635 (Loggerhead sea turtle (*Caretta caretta*) microbiome: insight into endozoic and epizoic communities - TurtleBIOME). The work of doctoral student KF has been fully supported by the “Young Researchers’ Career Development Project – Training of Doctoral Students” of the Croatian Science Foundation funded by the European Union from the European Social Fund.

## 11 Supplementary Material

**Figure S1**. Collection of epibiotic biofilm from skin (A) and carapace (B) of loggerhead sea turtles.

**Figure S2**. Principal coordinate analysis (PCoA) biplot of Bray-Curtis and weighted UniFrac distance for prokaryotic (16S, up) and eukaryotic diversity (18S, down); samples colored by factor “Sampling Locality”.

**Figure S3**. Principal component analysis (PCA) biplot of robust Aitchison distance for prokaryotic (16S, left) and eukaryotic diversity (18S, right); arrows indicate individual highly ranked ASVs that contribute to the displayed positions of the samples; lowest taxonomic assignment of each ASV is written in textboxes at the end of each arrow; samples colored by factor “Origin Sea”.

**Figure S4**. Principal coordinate analysis (PCoA) biplot of Bray-Curtis and weighted UniFrac distance for prokaryotic (16S, up) and eukaryotic diversity (18S, down) ; samples colored by factor “Origin Sea”.

**Figure S5**. Principal component analysis (PCA) biplot of robust Aitchison distance for prokaryotic (16S, left) and eukaryotic diversity (18S, right); arrows indicate individual highly ranked ASVs that contribute to the displayed positions of the samples; lowest taxonomic assignment of each ASV is written in textboxes at the end of each arrow; samples colored by factor “Season”.

**Figure S6**. Principal coordinate analysis (PCoA) biplot of Bray-Curtis and weighted UniFrac distance for prokaryotic (16S, up) and eukaryotic diversity (18S, down); samples colored by factor “Season”.

**Figure S7**. Principal component analysis (PCA) biplot of robust Aitchison distance for prokaryotic (16S, left) and eukaryotic diversity (18S, right); arrows indicate individual highly ranked ASVs that contribute to the displayed positions of the samples; lowest taxonomic assignment of each ASV is written in textboxes at the end of each arrow; samples colored by factor “Turtle State”.

**Figure S8**. Principal coordinate analysis (PCoA) biplot of Bray-Curtis and weighted UniFrac distance for prokaryotic (16S, up) and eukaryotic diversity (18S, down); samples colored by factor “Turtle State”.

**Figure S9**. Robust Aitchison PCA for prokaryotes / 16S rRNA gene (left) and eukaryotes / 18S rRNA gene (middle). Robust Aitchison PCA results were used for the procrustes analysis (right) as to compare prokaryotes (left) and eukaryotes (middle) samples positioning in the ordination spaces; the distances between each sample positions are indicated by a connecting line. Prokaryotic community samples are indicated by a full circle, while eukaryotic community samples are indicated by a full triangle. Significance (m2 value) and p-values were calculated by 999 series of MonteCarlo permutations.

**Table S1**. Sequencing results (raw counts) and taxonomy assignments of 16S dataset per ASV for all samples in this study.

**Table S2**. Sequencing results (raw counts) and taxonomy assignments of 16S dataset per ASV for all samples in this study.

**Table S3**. Sequencing results (raw counts) and taxonomy assignments of filtered cyanobacterial sequences of 16S dataset per ASV for all samples in this study.

**Table S4**. Relative abundance and absolute sequence count of prokaryotic families found on skin and carapace of loggerhead sea turtles.

**Table S5**. Core features of loggerhead sea turtles from 16S and 18S amplicon sequencing dataset.

## Notes

### Competing Interest Statement

The authors have declared no competing interest.

