## Supplemental_files for "Surface microbiota of Mediterranean loggerhead sea turtles unravelled by 16S and 18S amplicon sequencing": Kanjer-et-al-Supplements.docx

Supplementary Material

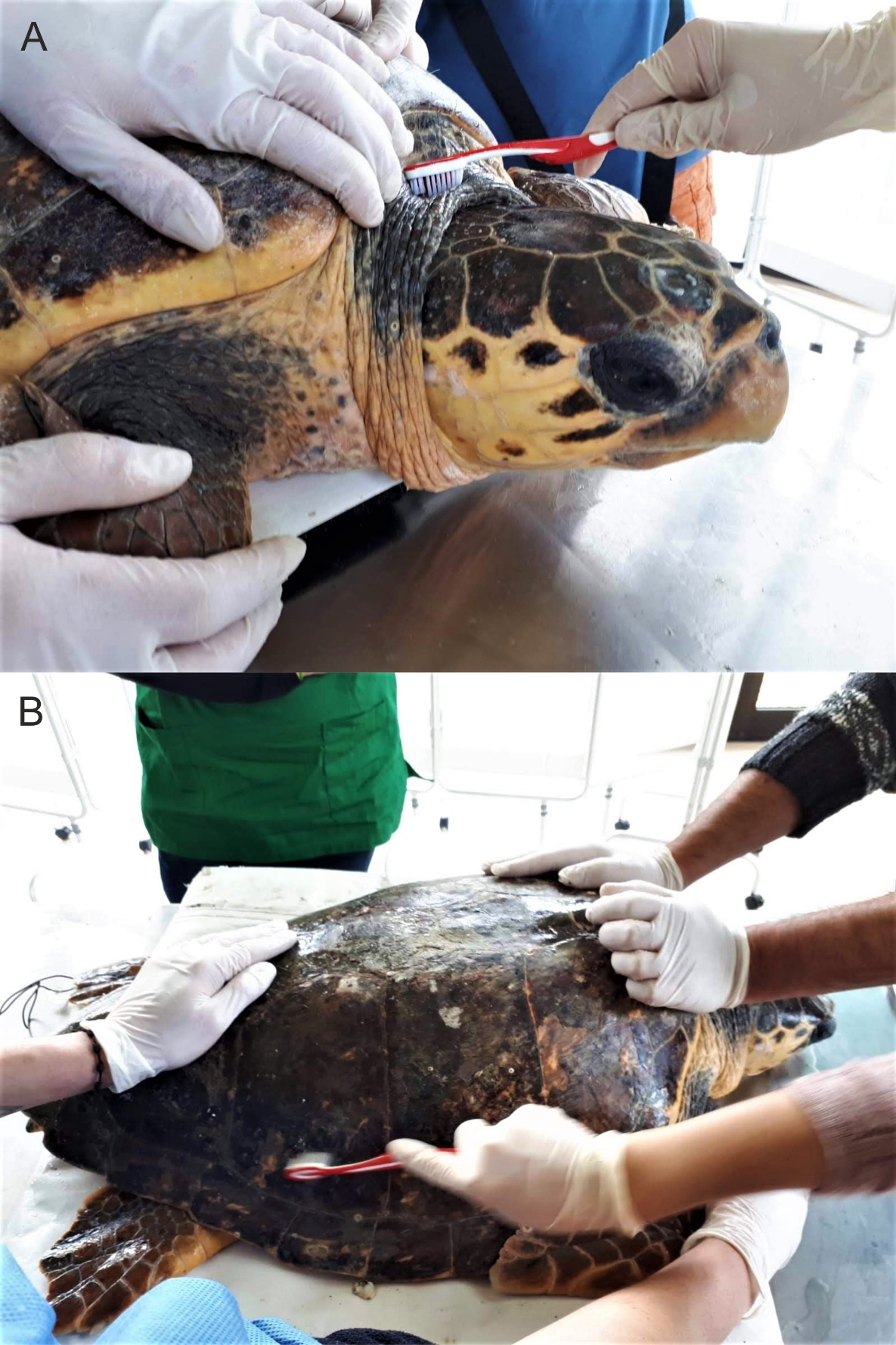

**Figure S1**. Collection of epibiotic biofilm scraping from skin (A) and carapace (B) of loggerhead sea turtles.

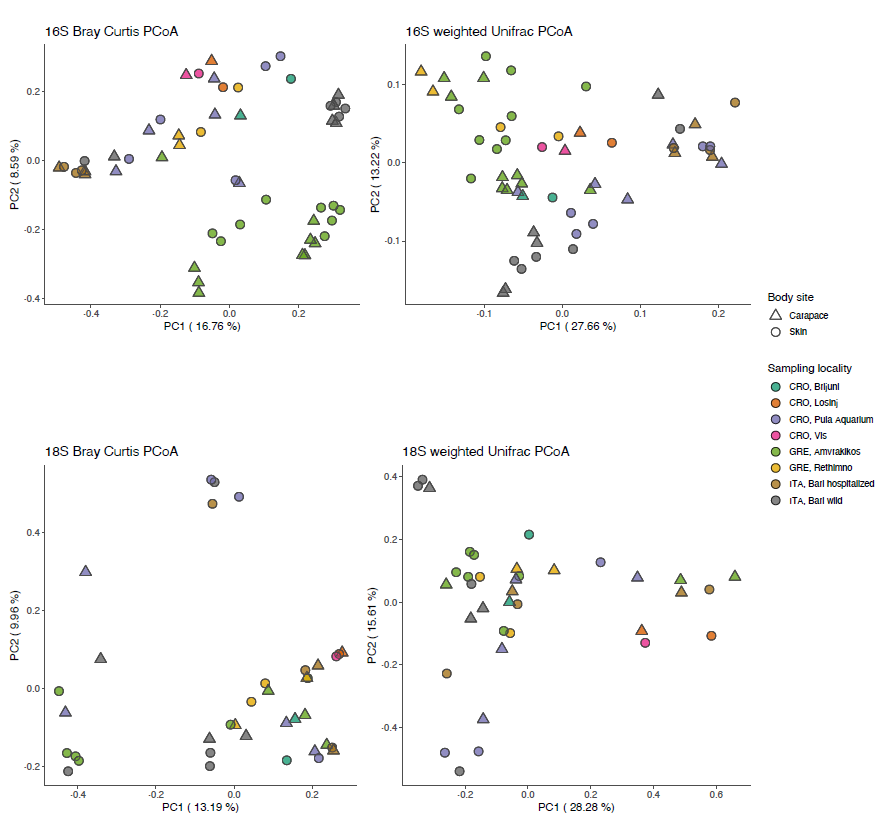

**Figure S2.** Principal coordinate analysis (PCoA) biplot of Bray-Curtis and weighted UniFrac distance for prokaryotic (16S, up) and eukaryotic diversity (18S, down); samples colored by factor “Sampling Locality”.

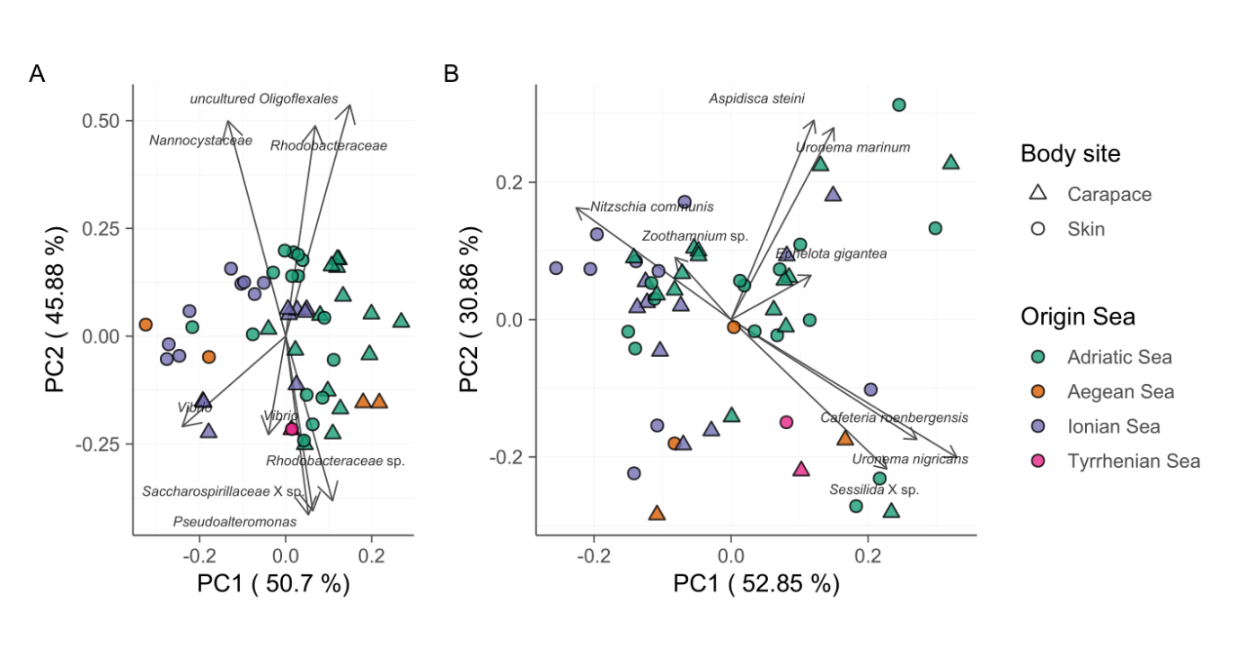

**Figure S3.** Principal component analysis (PCA) biplot of robust Aitchison distance for prokaryotic (16S, left) and eukaryotic diversity (18S, right); arrows indicate individual highly ranked ASVs that contribute to the displayed positions of the samples; lowest taxonomic assignment of each ASV is written in textboxes at the end of each arrow; samples colored by factor “Origin Sea”.

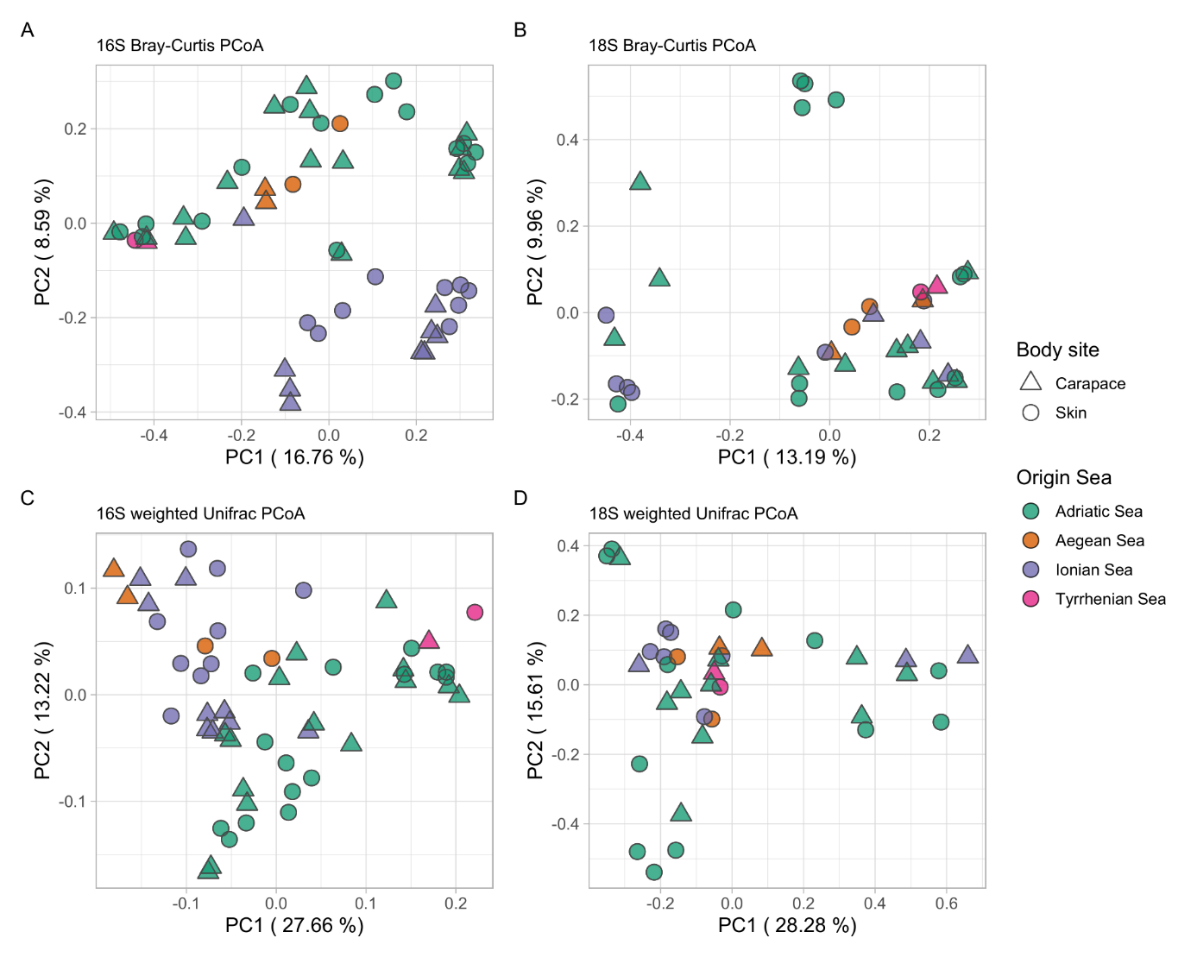

**Figure S4.** Principal coordinate analysis (PCoA) biplot of Bray-Curtis and weighted UniFrac distance for prokaryotic (16S, left) and eukaryotic diversity (18S, right); samples colored by factor “Origin Sea”.

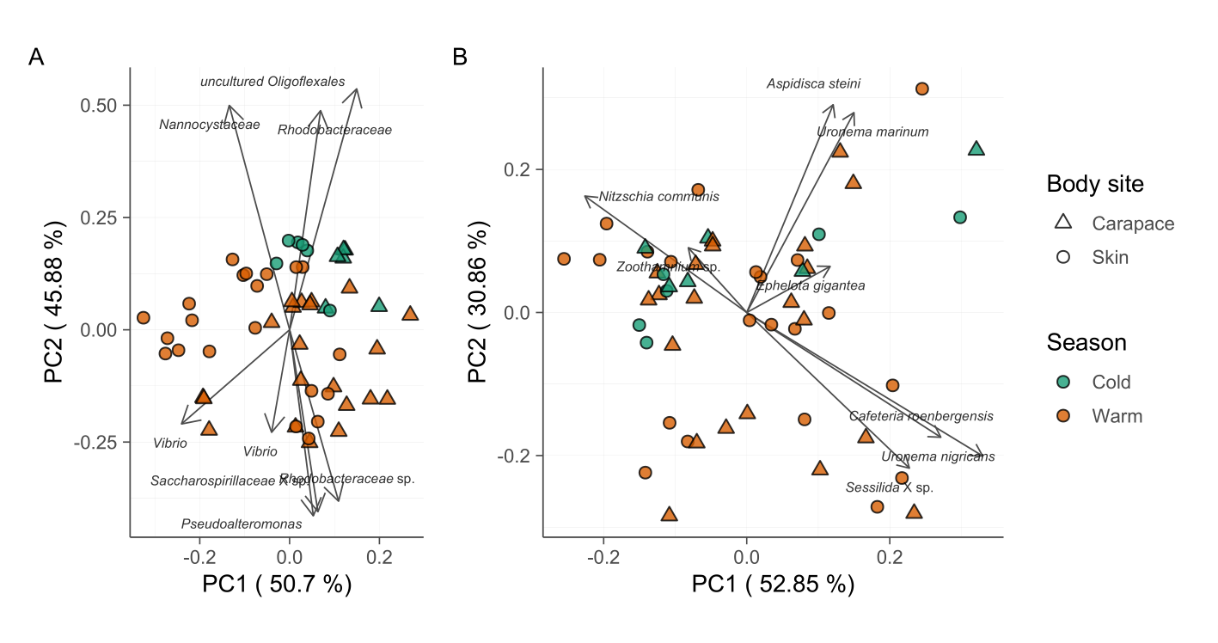

**Figure S5.** Principal component analysis (PCA) biplot of robust Aitchison distance for prokaryotic (16S, left) and eukaryotic diversity (18S, right); arrows indicate individual highly ranked ASVs that contribute to the displayed positions of the samples; lowest taxonomic assignment of each ASV is written in textboxes at the end of each arrow; samples colored by factor “Season”.

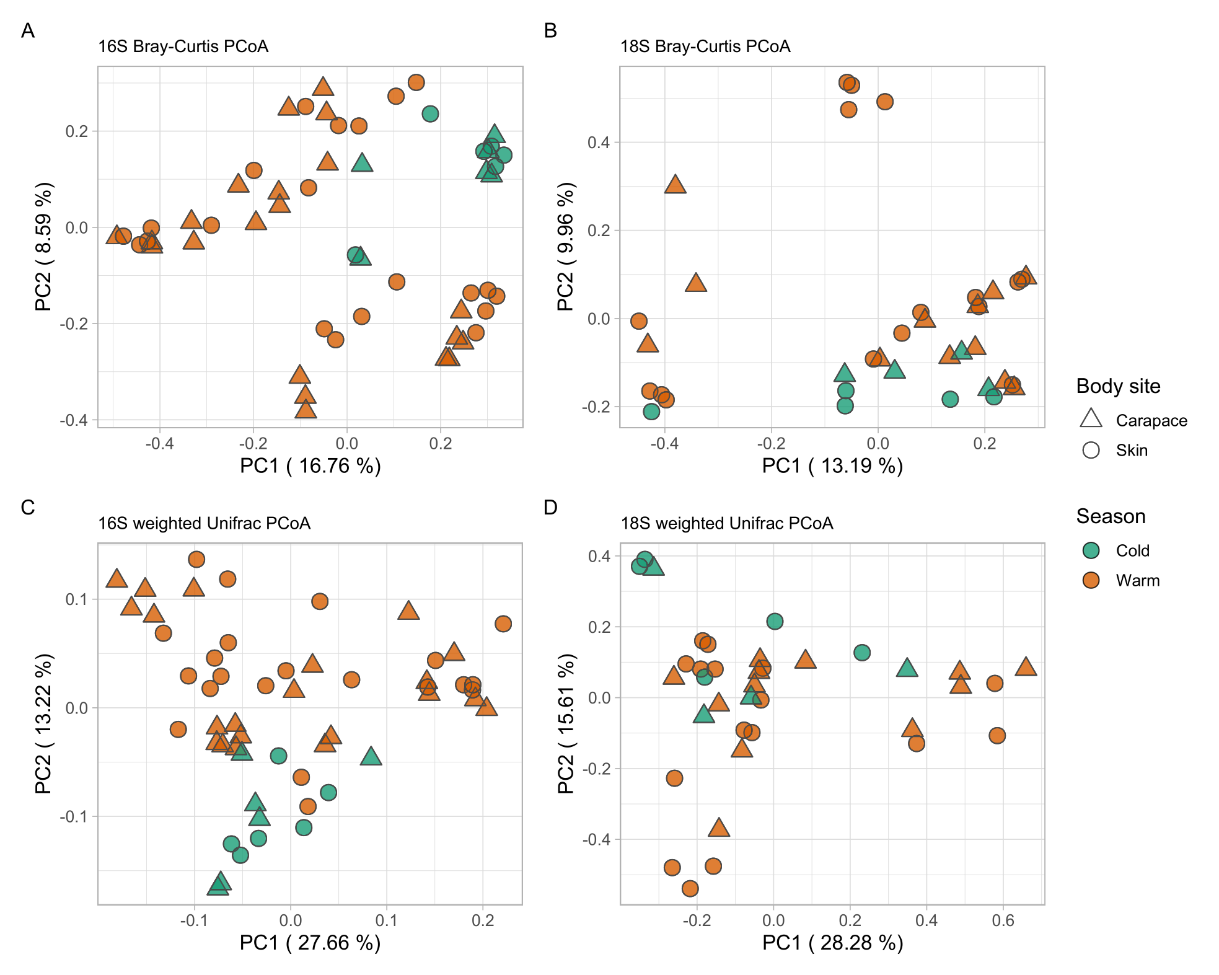

**Figure S6.** Principal coordinate analysis (PCoA) biplot of Bray-Curtis and weighted UniFrac distance for prokaryotic (16S, left) and eukaryotic diversity (18S, right); samples colored by factor “Season”.

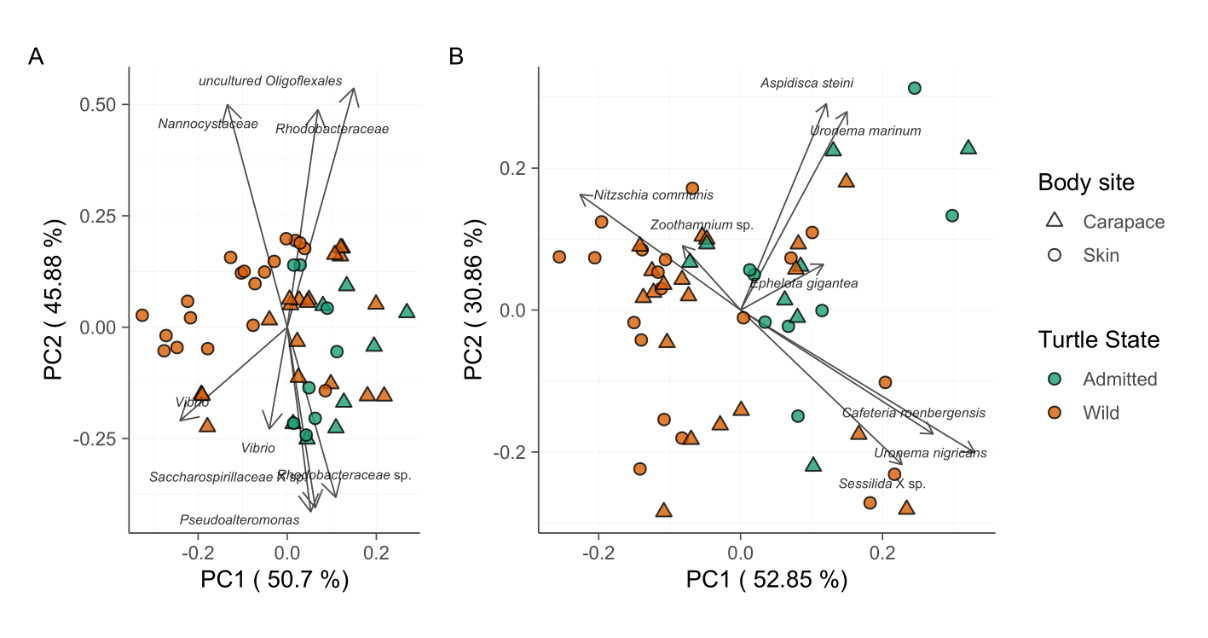

**Figure S7.** Principal component analysis (PCA) biplot of robust Aitchison distance for prokaryotic (16S, left) and eukaryotic diversity (18S, right); arrows indicate individual highly ranked ASVs that contribute to the displayed positions of the samples; lowest taxonomic assignment of each ASV is written in textboxes at the end of each arrow; samples colored by factor “Turtle State”.

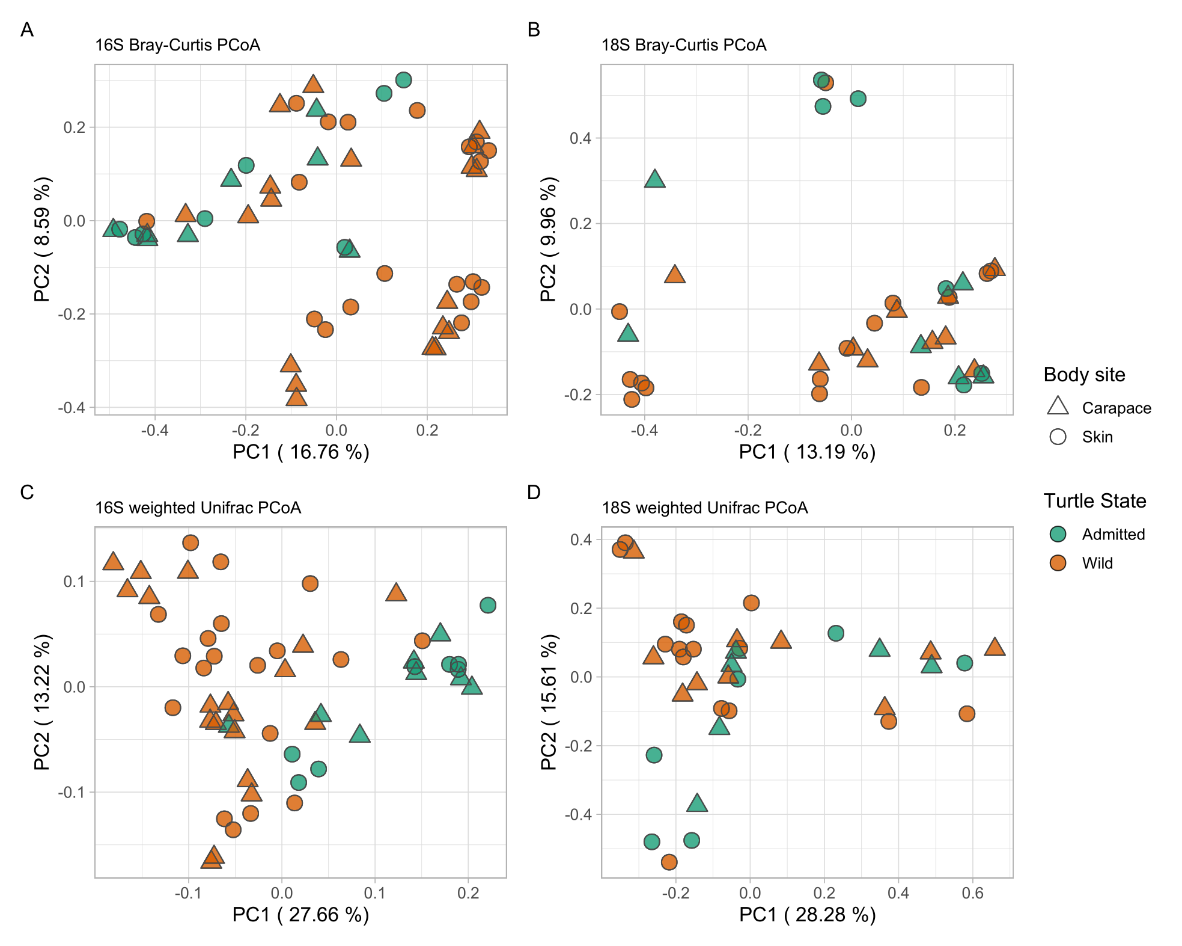

**Figure S8.** Principal coordinate analysis (PCoA) biplot of Bray-Curtis and weighted UniFrac distance for prokaryotic (16S, left) and eukaryotic diversity (18S, right); samples colored by factor “Turtle State”.

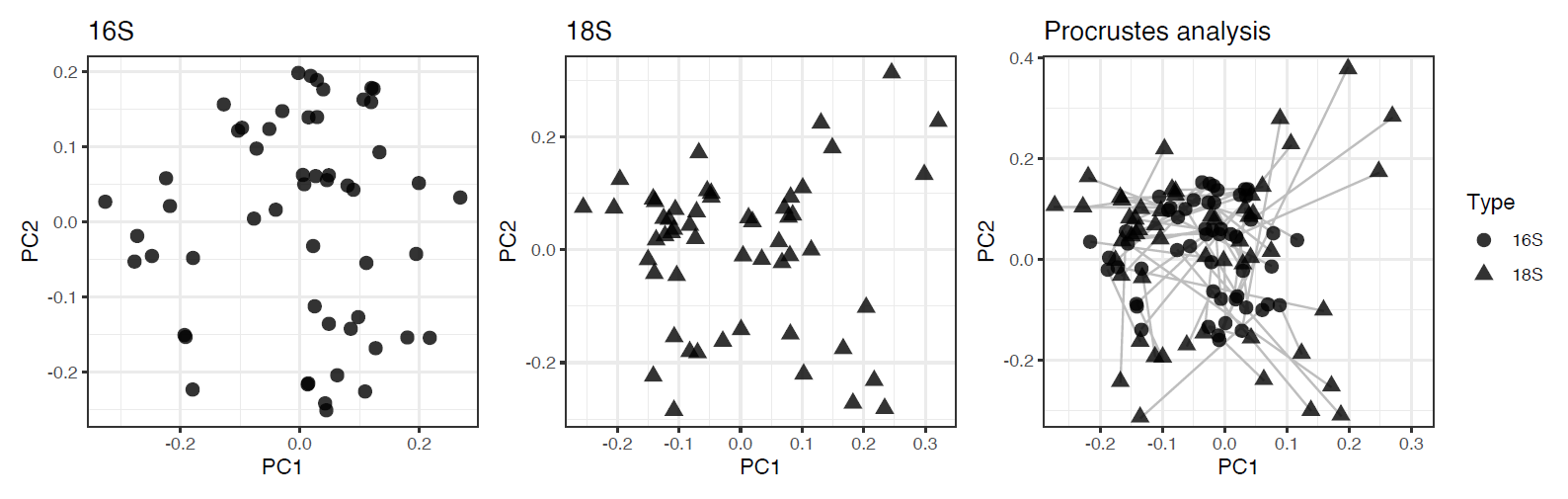

**Figure S9.** Robust Aitchison PCA for prokaryotes / 16S rRNA gene (left) and eukaryotes / 18S rRNA gene (middle). Robust Aitchison PCA results were used for the procrustes analysis (right) as to compare prokaryotes (left) and eukaryotes (middle) samples positioning in the ordination spaces; the distances between each sample positions are indicated by a connecting line. Prokaryotic community samples are indicated by a full circle, while eukaryotic community samples are indicated by a full triangle. Significance (m^2^ value) and p-values were calculated by 999 series of MonteCarlo permutations.
